# Narrow-window DIA: Ultra-fast quantitative analysis of comprehensive proteomes with high sequencing depth

**DOI:** 10.1101/2023.06.02.543374

**Authors:** Ulises H Guzman, Ana Martinez Del Val, Zilu Ye, Eugen Damoc, Tabiwang N. Arrey, Anna Pashkova, Eduard Denisov, Johannes Petzoldt, Amelia C. Peterson, Florian Harking, Ole Østergaard, Hamish Stewart, Yue Xuan, Daniel Hermanson, Christian Hock, Alexander Makarov, Vlad Zabrouskov, Jesper V. Olsen

**Affiliations:** Novo Nordisk Foundation Center for Protein Research, University of Copenhagen, Denmark; Thermo Fisher Scientific (Bremen) GmbH, Bremen, Germany; Thermo Fisher Scientific (San Jose), California, US

## Abstract

Mass spectrometry (MS)-based proteomics aims to characterize comprehensive proteomes in a fast and reproducible manner. Here, we present an ultra-fast scanning data-independent acquisition (DIA) strategy consisting on 2-Th precursor isolation windows, dissolving the differences between data-dependent and independent methods. This is achieved by pairing a Quadrupole Orbitrap mass spectrometer with the asymmetric track lossless (Astral) analyzer that provides >200 Hz MS/MS scanning speed, high resolving power and sensitivity, as well as low ppm-mass accuracy. Narrow-window DIA enables profiling of up to 100 full yeast proteomes per day, or ∼10,000 human proteins in half-an-hour. Moreover, multi-shot acquisition of fractionated samples allows comprehensive coverage of human proteomes in ∼3h, showing comparable depth to next-generation RNA sequencing and with 10x higher throughput compared to current state-of-the-art MS. High quantitative precision and accuracy is demonstrated with high peptide coverage in a 3-species proteome mixture, quantifying 14,000+ proteins in a single run in half-an-hour.

**Teaser:** Accurate and precise label-free quantification with comprehensive proteome coverage using narrow-window DIA

## Introduction

Genomics and proteomics have revolutionized our understanding of biology and hold tremendous potential for improving human health and the environment. In recent years, genomics has experienced a significant advancement with the introduction of next-generation sequencing (NGS), which has made DNA and RNA sequencing faster, more affordable, and more accurate. On the other hand, mass spectrometry (MS)-based proteomics has not witnessed a comparable leap in new technologies but has rather evolved incrementally through improvements in MS hardware, acquisition strategies, and associated software for analyzing MS data. Collectively, these advancements have transformed the way we study and analyze proteins and their modifications. State-of-the-art MS-based proteomics can comprehensively analyze the proteomes of human cell lines, covering the expression of virtually all proteins^1,2^. However, scalability remains a challenge, as achieving deep proteome coverage currently requires days of MS measurement time for each cell line^2,3^. This poses limitations for proteomics in comprehensive systems biology studies and large clinical cohort studies with a high number of patients, where coverage of low-abundance transcription factors and signaling proteins often needs to be sacrificed for throughput. In recent years, MS data acquisition methods have undergone a paradigm shift, with many applications transitioning from data-dependent acquisition (DDA) to data-independent acquisition (DIA)^4–6^. Faster and more sensitive instruments, coupled with advanced data processing software, have made DIA the preferred approach for maximizing proteome coverage in label-free single-shot analysis^7,8^. In this study, we present a single-shot analysis using DIA, which combines high-resolution MS1 scans with parallel, ultra-fast MS/MS scans of ∼ 200 Hz with high sensitivity. This empowers the use of narrow 2-Th DDA-like isolation windows for DIA, enabling the simultaneous comprehensive coverage of precursors afforded by DIA and the sensitivity and selectivity provided by narrow DDA-like isolation windows. We demonstrate that single-shot analysis facilitates comprehensive proteome profiling of complex organisms and is an ideal strategy for high-throughput proteomics profiling. For human proteome profiling, we introduce an optimized method that enables rapid and robust generation of ultra-deep proteomes (>12,000 proteins). This method combines offline high-pH reversed-phase (HpH) peptide fractionation with short 180-sample-per-day (SPD) online LC gradients and ultra-fast scanning DIA-MS, resulting in nearly complete coverage of the expressed human proteome within 4.5 hours of LC-MS/MS analysis.

## Results

### Orbitrap Astral MS enables narrow-window DIA with isolation width equivalent to DDA analysis

Clinical proteomics and systems biology studies require high throughput LC-MS/MS analyses with deep proteome coverage at high quantitative accuracy and precision. To achieve this, it is necessary to reduce MS instrument time usage by deploying shorter LC gradients with faster scanning MS instruments that can cope with the higher sample complexity per unit time. DIA has become the method of choice for single-shot deep proteome profiling with short gradients due to its high reproducibility and proteome coverage along with excellent quantitative performance. In DIA, co-eluting peptide ions are co-isolated and fragmented in predefined mass isolation windows, resulting in complex spectra containing fragment ions from multiple peptides that are analyzed together. Conversely, DDA has lower peptide sequencing capacity but higher specificity due to narrow isolation windows. In principle, higher specificity in DIA scans can be achieved by constraining the quadrupole isolation width similarly to DDA, but even state-of-the-art mass spectrometers cannot provide the sensitivity and acquisition speed needed to routinely perform DIA with 2-Th isolation windows on a chromatographic time-scale to maximize peptide sequencing across the mass range. To address this, we utilize a novel <u>as</u>ymmetric <u>tra</u>ck lossless (Astral) mass analyzer capable of ∼200Hz MS/MS acquisition rates at high resolving power and sensitivity, which permits routine DIA analysis with narrow 2-Th isolation windows. The analyzer is a part of the Thermo Scientific™ Orbitrap™ Astral™ mass spectrometer^9^ (Fig. 1a), which combines a modified Thermo Scientific™ Orbitrap Exploris™ 480^10^ mass spectrometer with the Astral analyzer via a transport octapole appended on the end of the Exploris ion routing multipole (IRM) and linking it to the ion processor (Fig. 1a). The function of the ion processor is similar to the combination of a C-Trap and IRM as it accumulates and fragments ions via higher-collisional dissociation (HCD)^11^ in a high-pressure region and then passes them to a lower-pressure region for extraction to the analyzer. The Astral analyzer represents an open electrostatic trap with pulsed detection^9^. While ions form nanosecond-wide packets undergoing temporal separation like in conventional orthogonal acceleration and/or multi-reflection time-of-flight (TOF) analyzers, the main differences are: (i) efficient transfer of ions from injection to detection with nearly lossless transmission, (ii) three-dimensional focusing of ions using the combination of grid-less asymmetric ion mirrors and an ion foil, (iii) significant spreading and overlapping of ion packets around the turning point of transversal drift to reduce space charge effects and (iv) high dynamic range detection that utilizes the conversion of accelerated ions into electrons that are converted into photons by a scintillator with subsequent detection by a photomultiplier.

**Figure 1.**
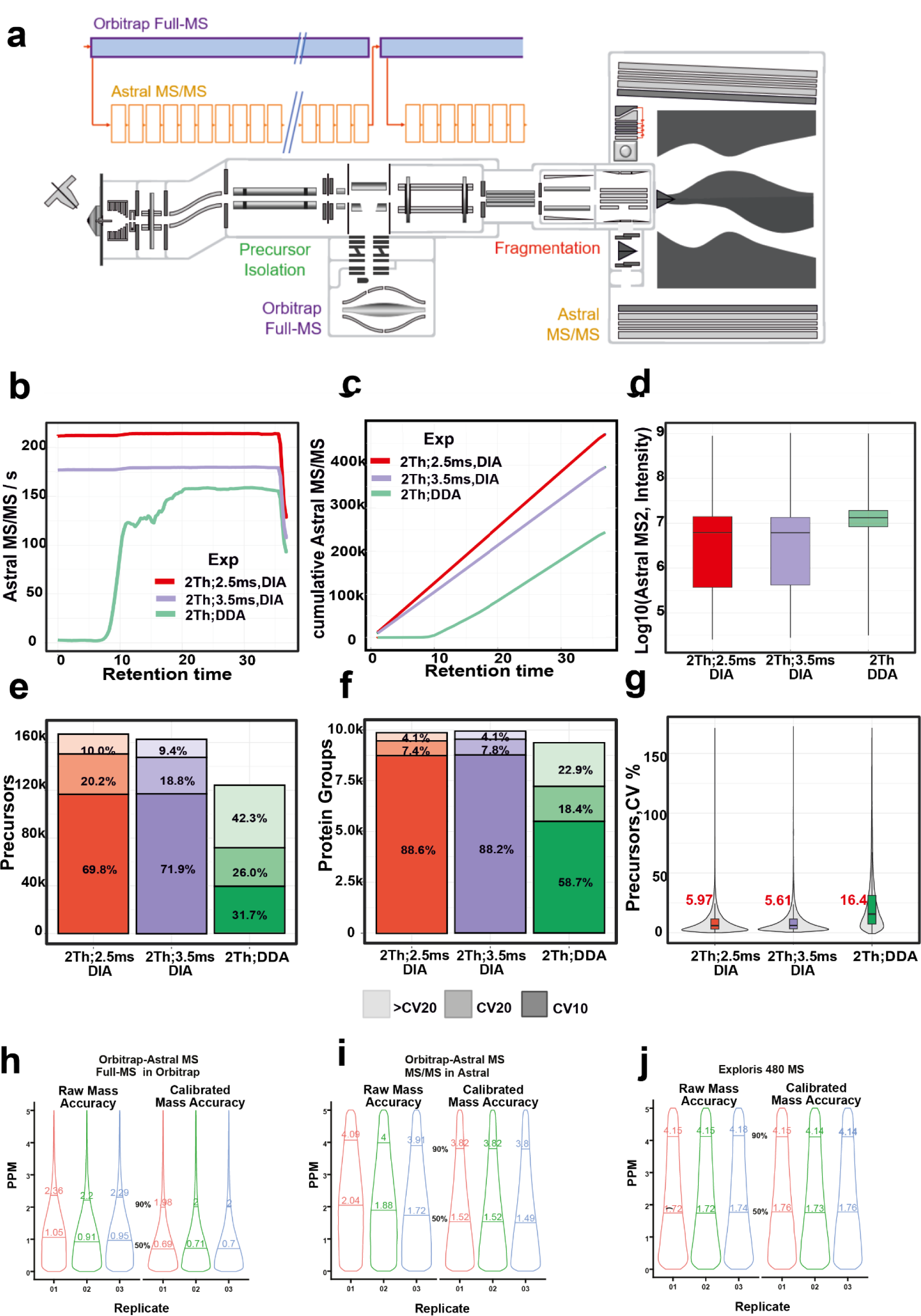
Benchmarking of narrow-window DIA against DDA. (a) Hardware overview of the Orbitrap Astral MS instrument. (b) MS/MS scan rate over the 28 min effective chromatographic gradient length. DDA and DIA slow (3.5 ms) and fast (2.5 ms) methods are depicted by different colors (green, purple and red respectively) (c) Cumulative MS/MS scans across 28 min effective chromatographic gradient acquired in Astral analyzer. (d) Log 10 intensity of MS/MS spectra measured in the Astral analyzer with DDA and DIA (e) Quantified precursors measured with DDA and narrow-window DIA operation mode below 10 and 20 % of coefficient of variation (CV) (n=3, 200 ng tryptic HEK293 peptides). (f) Quantified and identified proteins below 10 and 20 % of coefficient of variation (CV) with DDA and narrow-window DIA approaches (n=3, 200 ng tryptic HEK293 peptides). (g) Coefficients of variation (CVs, %) for quantified precursors by different operation modes. Median CV is shown in red for each approach. (h-j) absolute median mass deviation of peptide precursors determined in Orbitrap Astral Full-MS using the Orbitrap analyzer (f), Orbitrap Astral MS/MS using the Astral analyzer (h) and Orbitrap Exploris 480 MS using the Orbitrap analyzer (j).

Ion injection from the trapping ion processor into the analyzer is synchronized to maximize scan duty cycle and the unique transversal asymmetric oscillations combine temporal and spatial focusing with an asymmetric ion track length of 30 meters resulting in high resolution of >80,000 FWHM at *m/z* 524. The nearly lossless ion transfer due to optimized ion refocusing during ion injection, oscillations, and high dynamic range detection enable high MS/MS scan rates of ∼200 Hz at very high sensitivity without the need for averaging scans. Full scans can be analyzed in parallel at ultra-high resolution and dynamic range with the Orbitrap mass analyzer. Automatic Gain Control (AGC) and a high dynamic range detector enable high inter-scan dynamic range. It is possible to operate the instrument in DDA mode with top speed acquisition of ∼150 Hz, but restricting the maximum ion fill time to 3.5 ms in DIA mode results in 170 Hz MS/MS acquisition rate while 2.5 ms ion injection time delivers >200Hz MS/MS acquisition with 2-Th isolation windows (Fig. 1b). The fastest DIA scanning speed generates ∼400,000 MS/MS spectra in half an hour of LC-MS/MS analysis of a human embryonic kidney cell (HEK 293) tryptic digest (Fig. 1c). Even though, the overall MS/MS spectra intensity was higher in DDA mode (Fig. 1d), DIA acquisition mode results in higher identifications of approximately 170,000 peptide precursors and ∼10,000 proteins groups (hereafter referred as proteins) (Fig. 1e,f). Importantly, the quantitation of peptides and protein identifications can be performed with high reproducibility with coefficient of variation (CV) below 20% for 90% of precursors and 95% of proteins, respectively (Fig. 1e,f). The median CVs at precursor level for DIA approaches were lower compared to DDA (< 6% vs < 17% respectively) (Fig. 1g). This observation can be explained by the stochastic nature of DDA precursor selection^12^. Comparison of DDA and DIA strategies showed higher performance for DIA, although closely followed by DDA.

Fragment ion mass accuracy is an important parameter for determining fragment ion identity and restricting matches in peptide search engines^13^. To evaluate the mass accuracy provided by the Astral mass analyzer in MS/MS mode, we analyzed a tryptic digest of a whole cell lysate of HEK 293 cells using a 200Hz 2-Th DIA method. The absolute median mass deviation of the peptide precursors determined in the Orbitrap MS1 full-scans recorded at 240,000 resolution was 0.95 ppm (Fig. 1h), which is comparable to previous reports^14^. Analyzing all matched fragment ions recorded with the Astral analyzer, we found that the median of absolute mass deviation to be 1.88 ppm (Fig. 1i). This is on par with Orbitrap MS/MS measurements at 15,000 resolution (Fig. 1j). While Orbitrap mass calibration is stable for days, the Astral mass accuracy may drift over time with a mass error of ∼3 ppm. Importantly, this slight drift in mass measurement accuracy in the Astral analyzer over time can readily be corrected using a regular automatic recalibration via the internal calibrant source. Alternatively, it can also be recalibrated post-acquisition, leading to a sub-ppm level mass accuracy in most recalibrated masses at both MS1 and MS2 levels.

### Fast sequencing enables profiling of comprehensive proteomes with short LC gradients

Complete proteome coverage is important for large-scale systems biology studies with hundreds or thousands of conditions. The first complete proteome of a eukaryotic organism to be completed by mass spectrometry was yeast^15^ and with state-of-the-art instrumentation it is possible to cover ∼4000 yeast proteins in approximately 1 h of LC-MS/MS time^16^. To assess how much time is needed to measure the expressed yeast proteome (∼4500 proteins) with a fast-scanning narrow-window DIA strategy (3-Th), we analyzed whole yeast cell tryptic digests using different LC gradient lengths from 5-min LC gradients (8-min from injection-to-injection, equivalent to 180 SPD) up to a 96 SPD (15-min). Remarkably, close to complete yeast proteome coverage was achieved with all gradients highlighting the possibility to analyze the yeast proteome ten times faster than previously reported with higher coverage and high quantitative reproducibility (Fig. 2a). The average yeast protein sequence coverage achieved with 180 SPD was 25% which was improved to 30% with the 12-min and 15-min gradients (Fig. 2b).

**Figure 2.**
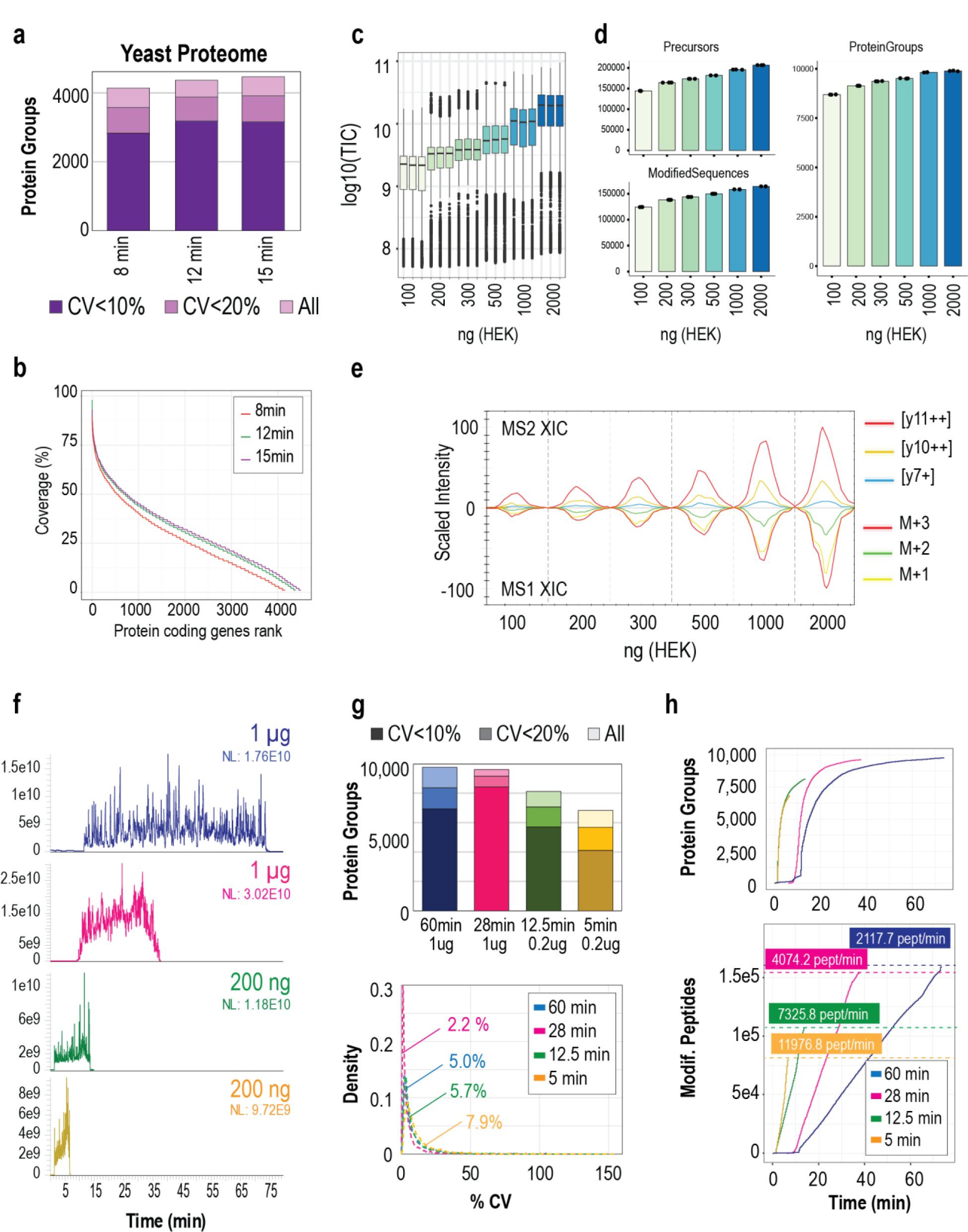
Deep single-shot proteomes with variable LC gradients. (a) Protein groups identified and quantified below 10 and 20% of coefficient of variation (n=3) in a yeast extract using different chromatographic gradient lengths. (b) Proteins ranked by sequence coverage in different chromatographic gradient lengths. (c) Boxplot of Total Ion Current (in log10 scale) measured in a dilution series (from 100 to 2000 ng) of a HEK293 cells protein digest. (d) Protein groups, precursors and modified peptides identified (n=3) in a dilution series of HEK293 cells peptide extract using Spectronaut (v17). (e) Extracted ion chromatogram (top MS2, bottom MS1) of peptide DYFGAHTYEKKAK(3+) in a dilution series of HEK293 peptides, from 100 to 2000 ng. (f) Total ion chromatograms of different MS runs of HEK293 peptide extracts. From top to bottom: 60 minutes and 1µg load, 28 minutes and 1 µg load, 12.5 minutes and 200 ng load and 5 minutes and 200 ng load. (g) Protein Groups (top) identified and quantified below 10 and 20% of coefficient of variation (n=3) in different loadings and gradients of HEK293 cells peptide extract using Spectronaut (v18). Below, distribution of protein groups quantitation coefficient of variation for each gradient. Number in the plot indicates the median CV. (h) Cumulative identifications of protein groups (top) or modified peptides (bottom) for 1 µg in 28 or 60 minutes of chromatographic separation, or 200 ng in 12.5 or 5 minutes of chromatographic separation.

The human proteome is more complex than the yeast proteome comprising more than 12,000^1,17^ protein coding genes with a higher dynamic range (4 to 6-log)^18,19^. Consequently, comprehensive coverage of a human cell line proteome requires longer MS measurement time. To evaluate the human proteome coverage and sensitivity achieved with narrow-window DIA (2-Th), we measured increasing amounts of HEK 293 protein digests (from 100ng to 2000ng) with 60-min active LC gradients. Reassuringly, the measured MS/MS signal scaled with loading amounts and did not seem to be saturated at the highest load (Fig. 2c). Peptide and protein coverage and quantitative reproducibility also correlated with loading amounts with an average of 9,879 proteins and 159,627 unique peptides (164,216 modified peptides) when injecting 2-µg HEK 293 protein digest, but, importantly, we could cover 88% of the proteins even after reducing the input to 100-ng (Fig. 2d). Similar quantitative performance was observed between full-scan level with Orbitrap analyzer measurements and MS/MS level using the Astral analyzer as judged by the extracted ion chromatogram of a peptide precursor and its highest fragments across the different loading amounts (Fig. 2e). To find the best compromise between proteome coverage and invested MS measurement time, we compared different loadings of the HEK 293 digest using different active LC gradient lengths of 5-min, 12.5-min, 28-min and 60-min (Table 1 and 2) (Fig. 2f). Interestingly, the 28-min gradient analysis of 1-µg resulted in 9,619 proteins covered with outstanding reproducibility and a median CV below 3 % (Fig. 2g). This better performance could be explained by the increased scanning speed used for those runs, in which we employed 2.5 ms of injection time and achieved >200 Hz of DIA-MS/MS scanning speed. Using shorter gradients (28 min vs 60 min) for the same injection load boosted the MS signal (Fig. 2f) and allowed us to reduce the injection time from 3.5 to 2.5 ms without losing sensitivity. This enables an increased scanning speed, resulting in similar protein and peptide coverage in half the acquisition time (Fig. 2g-h). Notably, using 200-ng and 12.5-min gradient we detected up to 8120 proteins with median CVs below 6% (Fig. 2g). Most importantly, same loading with a 5-min gradient reached 6828 proteins (Fig. 2g), 3500 of which were covered within the first minute of peptide elution (Fig. 2h). This was due to the very high identification rate of ∼11,000 peptides/min and >1000 proteins/min, meaning even narrow window 2-Th DIA scans result in chimeric spectra from the Astral analyzer that can be deconvoluted with software strategies with faster gradients and high loads (Fig. 2h).

**Table 1:** LC parameters overview for the different experiments presented in the text

| Experiment | Input | Gradient<br>(peptide elution<br>duration) | Total<br>acquisition<br>time | SPD | Column |
| --- | --- | --- | --- | --- | --- |
| DDA vs DIA comparison.<br>DDA mode. | HEK293, 1000 ng | 28 min | 37 min | 36 | μPAC column (110 cm) |
| DDA vs DIA comparison.<br>DIA mode 2.5 ms. | HEK293, 1000 ng | 28 min | 37 min | 36 | μPAC column (110 cm) |
| DDA vs DIA comparison.<br>DIA mode 3.5 ms. | HEK293, 1000 ng | 28 min | 37 min | 36 | μPAC column (110 cm) |
| Yeast proteome | *S.cerevisiae* 200ng | 5 min | 7 min | 180 | PepMap (2 μm, 150 μm x<br>15 cm) |
| Yeast proteome | *S.cerevisiae* 200ng | 9 min | 12 min | 120 | PepMap (2 μm, 150 μm x<br>15 cm) |
| Yeast proteome | *S.cerevisiae* 200ng | 12 min | 15 min | 96 | PepMap (2 μm, 150 μm x<br>15 cm) |
| Single-shot | HEK293, 1000 ng | 60 min | 80 min | 13 | PepMap (2 μm, 150 μm x<br>75 cm) |
| Single-shot | HEK293, 1000 ng | 28 min | 38 min | 36 | μPAC column (110 cm) |
| Single-shot | HEK293, 200 ng | 12.5 min | 15 min | 96 | PepMap (2 μm, 150 μm x<br>15 cm) |
| Single-shot | HEK293, 200 ng | 5 min | 7 min | 180 | PepMap (2 μm, 150 μm x<br>15 cm) |
| Dilution series | HEK293 (from 100 to<br>2000 ng) | 60 min | 80 min | 13 | PepMap (2 μm, 150 μm x<br>75 cm) |
| Window Optimization.<br>2Th. | HeLa (10, 25 and 100<br>ng) | 60 min | 80 min | 13 | PepMap (2 μm, 150 μm x<br>75 cm) |
| Window Optimization.<br>4Th. | HeLa (10, 25 and 100<br>ng) | 60 min | 80 min | 13 | PepMap (2 μm, 150 μm x<br>75 cm) |
| Window Optimization.<br>8Th. | HeLa (10, 25 and 100<br>ng) | 60 min | 80 min | 13 | PepMap (2 μm, 150 μm x<br>75 cm) |
| Window Optimization.<br>16Th. | HeLa (10, 25 and 100<br>ng) | 60 min | 80 min | 13 | PepMap (2 μm, 150 μm x<br>75 cm) |
| Quantification Benchmark | *E.coli*, *H.sapiens* and<br>*S. cerevisiae*<br>(variable mixtures) | 28 min | 37 min | 36 | μPAC column (110 cm) |
| Fractionation | HEK293 200<br>ng/fraction | 5 min | 7 min | 180 | PepMap (2 μm, 150 μm x<br>15 cm) |
| Yeast KO | *S.cerevisiae*, 200ng | 5 min | 7 min | 180 | PepMap (2 μm, 150 μm x 15 cm) |
| Clinical samples | Human tissue, 500 ng | 5 min | 7 min | 180 | PepMap (2 μm, 150 μm x 15 cm) |

**Table 2:**
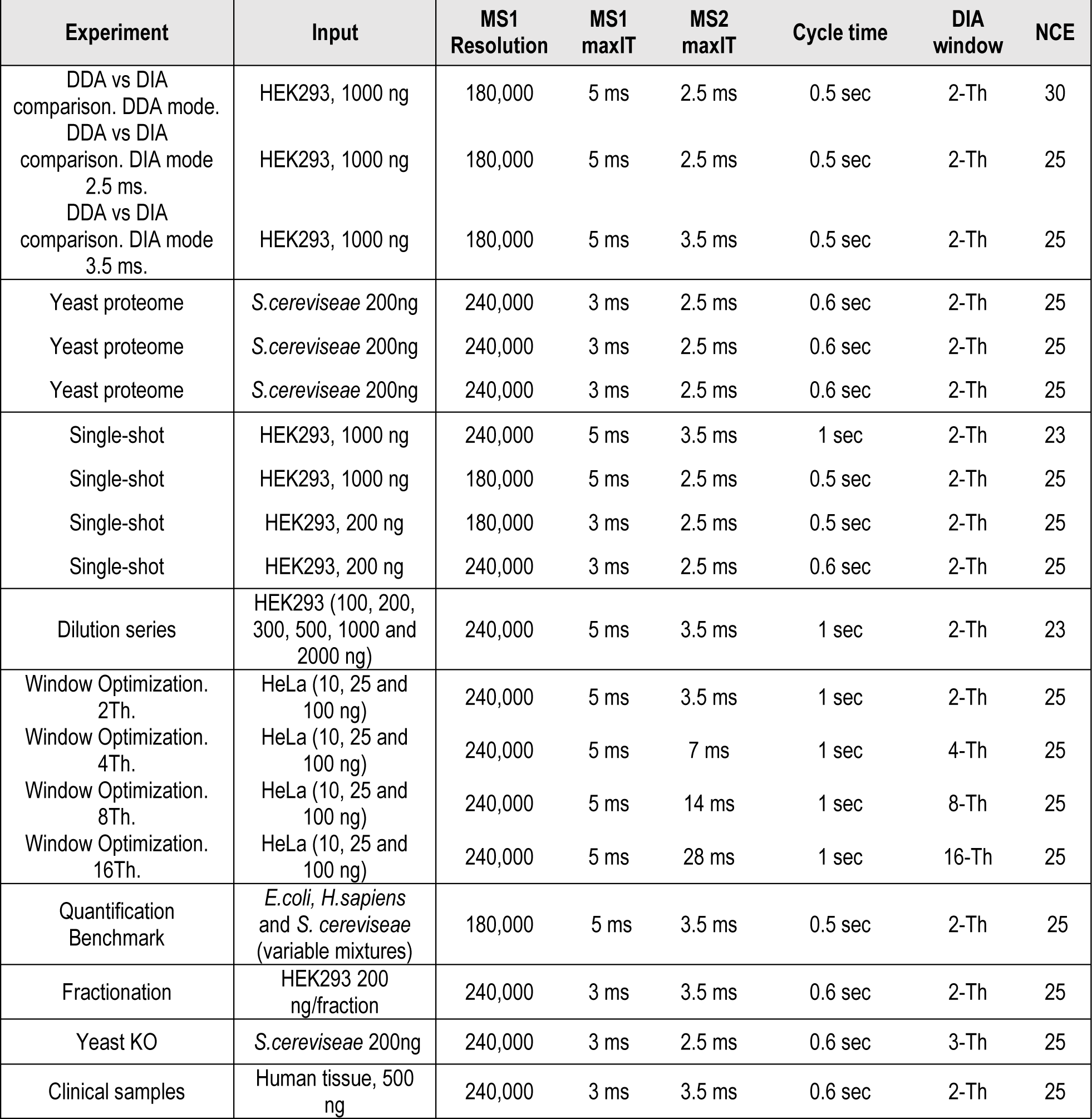
LC parameters overview for the different experiments presented in the text

### Systematic optimization of window placement as a function of precursor mass-to-charge

The observed reduction in identifications with lower sample loads indicated a need for tailored DIA acquisition methods, especially for high-sensitivity strategies. We tested the impact of increasing the isolation window width and scaling the maximum injection time (maxIT) accordingly to decreasing sample amounts, while keeping scan cycle time constant (Fig. 3a). We analyzed 10-ng, 25-ng and 100-ng injections using 2-Th, 4-Th, 8-Th and 16-Th windows with 3.5-ms, 7-ms, 14-ms and 28-ms maxIT (Table 1 and 2), respectively, using an 80 minutes acquisition (corresponding to 60 minutes of peptide elution). Interestingly, we found that 4-Th windows with 7-ms maxIT was the optimal method for 100-ng injections, resulting in 8,600 protein, whereas lower loads benefited from widening the isolation window to 8-Th, resulting in 6,600 proteins from just 10-ng of protein digest input (Fig. 3b). With decreasing loading amounts, the ideal peptide ion target value is rarely reached and the under-sampling is most pronounced with narrow isolation windows (Fig. 3c). The impact of the different window sizes was not uniform across the mass range with peptides of higher *m/z* most affected (Fig. 3d). Therefore, scaling window size and maxIT in different mass ranges according to sample loads may provide a means to set up the most optimal method for maximizing peptide coverage given the sample input.

**Figure 3.**
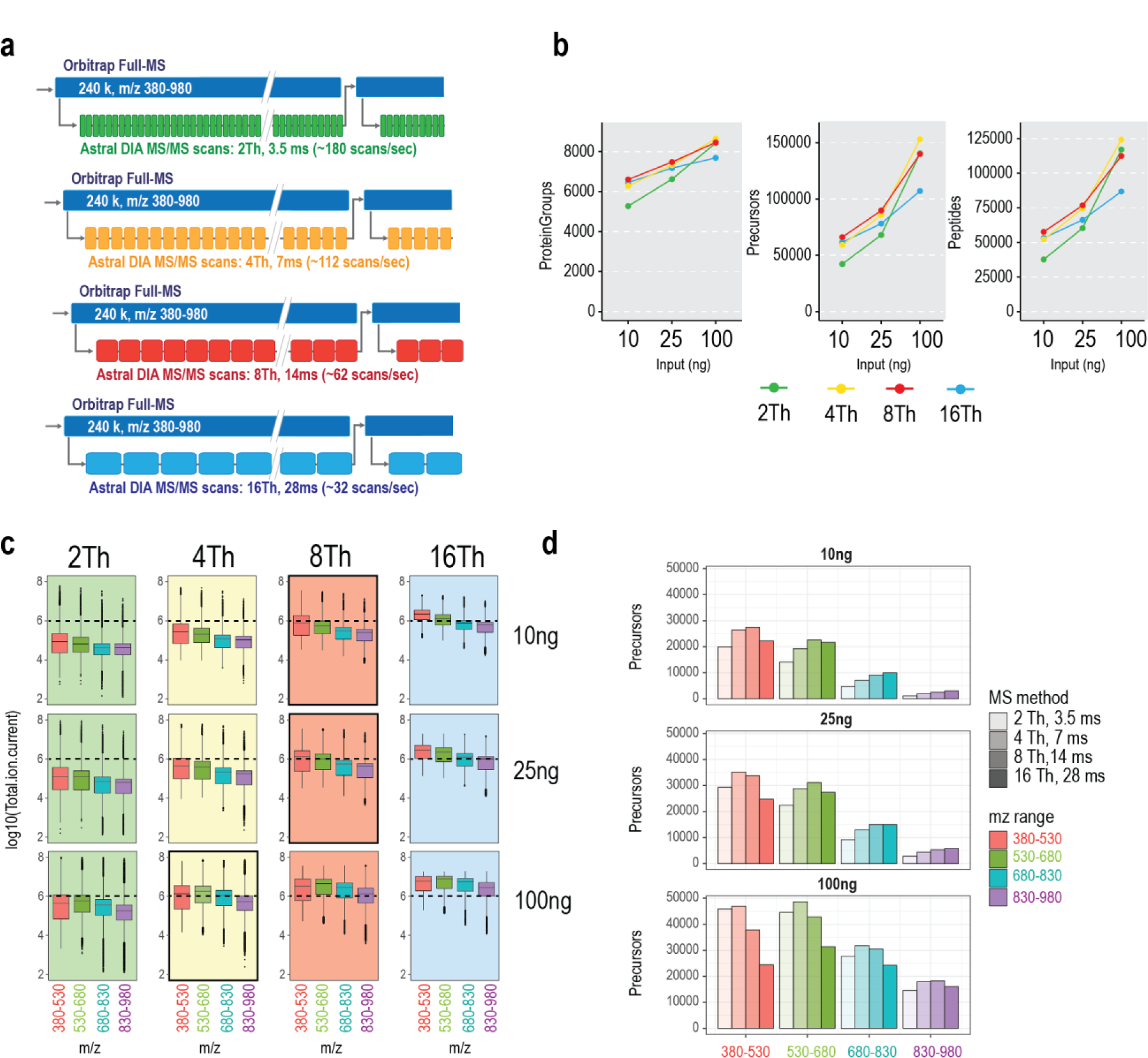
Optimization of DIA acquisition parameters for low peptide loads. (a) Schematic representation of acquisition cycles in the Orbitrap-Astral analyzers with different maxIT and window width parameters. Orbitrap MS1 scans were acquired at 240K while in parallel Astral scans were acquired during a duty cycle of 1 second. For each method, the injection time was scaled to the window width, so the same mass range is covered in each cycle: 2Th and 3.5ms, 4Th and 7ms, 8Th and 14ms, 16Th and 28ms. (b) Protein groups, precursors and peptides (average of n=3) identified for 10, 25 and 100 ng using different window widths, as depicted in panel A. (c) Boxplots of total ion current measured at MS2 level, grouped in four mass ranges (380-530, 530-680, 680-830, 830-980), for different sample loadings and different acquisition methods. Highlighted, the method that gives higher identifications for that sample load. Black dashed line indicates the highest median intensity obtained for 100 ng and 4Th, which gives the highest identification rate. (d) Precursors (average of 3 technical replicates) identified in four mass ranges (380-530, 530-680, 680-830, 830-980) using different acquisition methods (injections times and window width).

### Enhanced quantitative precision and accuracy for label-free quantification

Next, to evaluate the quantitative performance of the narrow window DIA methods for label-free quantification (LFQ), we analyzed mixed species proteome samples composed of tryptic digests of human, yeast, and *E.coli* proteins mixed in six specific ratios (Fig. 4a). We analyzed the samples with narrow-window DIA (Table1 and 2) and short LC gradients in order to benchmark the proteome coverage and quantitative performance of the Orbitrap Astral MS in terms of accuracy and precision against an Orbitrap Exploris 480 MS. The Orbitrap Astral MS was able to identify >14,000 proteins and >260,000 peptide precursors in a 28-min LC gradient when analyzing 800-ng on column using Spectronaut (v17) (Fig. 4b). In comparison, an Orbitrap Exploris 480 MS identified ∼7,000 proteins and ∼76,000 peptide precursors in a 45-min LC gradient (Table 1 and 2), demonstrating that the Orbitrap Astral MS identifies >2x proteins and 3.5x peptide precursors in shorter analysis time. Notably, ∼90% of proteins had CVs of less than 20% and the number of missing values in technical triplicates were very low (<3%), indicating high technical reproducibility. At all three expected ratios (1.5x, 4x, and 9x) for yeast and *E.coli*, MS/MS-based protein quantification results in Astral analyzer were more accurate with lower standard deviations (∼0.3) compared to the Orbitrap Exploris 480 MS (Fig. 4c). The quantification accuracy of precursors at MS2 level was as good as the Orbitrap Exploris 480 MS, indicating that the Astral analyzer is as quantitative as the Orbitrap analyzer. Additionally, the significantly higher protein coverage and number of peptide precursors measured per protein resulted in superior protein quantification, particularly when using the MaxLFQ algorithm^20^. Compared to the QUANT 2.0 algorithm that only uses the top 3 peptide groups for protein quantification, almost all the measured (>99%) precursors were utilized for protein quantification in the MaxLFQ algorithm (Fig. 4d). Notably, the additional ∼210,000 precursors disregarded by the QUANT 2.0 algorithm were almost as abundant overall as the ∼44,000 that were selected^21^ (Figure 4e). When incorporated in the MaxLFQ algorithm, these 5x extra precursors led to much improved quantification performance. This demonstrates the advantages of the narrow window DIA strategy for accurate and precise protein quantification.

**Figure 4.**
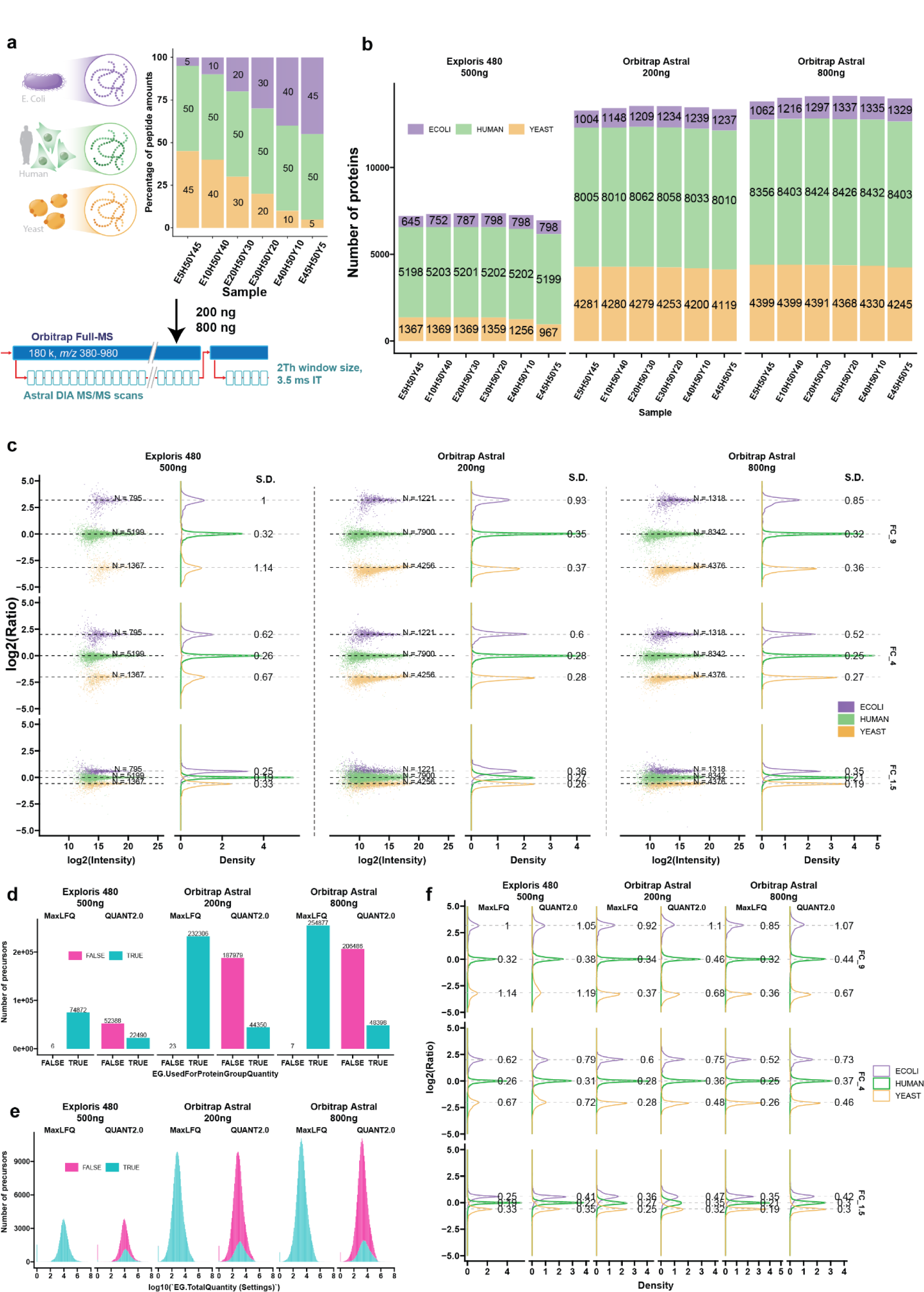
Enhanced LFQ Accuracy and Precision with Orbitrap Astral MS. (a) Graphical representation of the experimental design. Tryptic peptides from three species were combined in six distinct ratios (E5H50Y45, E10H50Y40, E20H50Y30, E30H50Y20, E40H50Y10, and E45H50Y5). Samples were processed using Orbitrap Astral MS in technical triplicates, employing a 3.5 ms maxIT and 2-Th window size method. The loading amounts were 200-ng and 800-ng. (b) Number of proteins identified from the three species in each sample. For the Orbitrap Exploris 480 MS runs, the loading amount was 500-ng. (c) Log-transformed ratios of quantified proteins. Scatter plots for all runs over the log-transformed protein intensities are displayed on the left, while density plots are on the right. Colored dashed lines represent expected log2(A/B) values for proteins from humans (green), yeast (orange), and E. coli (purple). Standard deviations are displayed on the density plots. FC_9, FC_4, and FC_1.5 are calculated from log2(E45H50Y5/E5H50Y10), log2(E40H50Y10/E10H50Y40), and log2(E30H50Y20/E20H50Y30), respectively. (d) Number of precursors used and not used for protein quantifications in the MaxLFQ and QUANT 2.0 algorithms. (e) The intensity distribution of precursors used for protein quantifications in MaxLFQ and QUANT 2.0 algorithms. (f) Density plots of log-transformed protein ratios quantified using both the MaxLFQ and QUANT 2.0 algorithms.

### Fast multi-shot acquisition strategy to obtain deep proteomes

The main obstacle to overcome for complete determination of human proteomes is the inherent wide dynamic range of individual protein abundances. A multi-shot proteomics strategy increases the dynamic range and coverage compared with single-shot experiments in human proteome investigations by decreasing the complexity of the sample introduced into the LC-MS, but usually comes at the cost of extensive measurement time. To maximize proteome coverage, we took advantage of high-resolution offline high pH reversed-phase peptide chromatography (HpH)^22,23^ in combination with short online LC gradients and narrow-window DIA (2-Th) analysis on the Orbitrap Astral MS performing >10,000 high-resolution MS/MS scans per minute. To identify the most effective strategy to comprehensively cover a human cell line proteome, we fractionated a tryptic digest of HEK-293 cells into 46, 34, 23 or 12 HpH-fractions (Fig. 5a). Analyzing 200-ng from each of the 46 fractions with 5-min effective gradient DIA runs constituting a total of 6 h measurement time resulted in the identification of 12,179 protein-coding genes and 222,389 peptide sequences using Spectronaut (v17) with directDIA+. These results represent a 6-fold decrease of required MS acquisition time and 5-fold reduction of sample input needed as compared to prior approaches (Fig. 5a). Importantly, high quantitative performance was maintained between different fractionation schemes with Pearson correlation coefficients above 0.9 for all pairwise comparisons (Fig. 5b). Combining three biological replicates of each of the fractionation schemes resulted in a maximum coverage of 12,328 protein-coding genes from ∼246,000 peptides from both 34 and 46 fractions (Fig. 5c). Notably, the 34 and 46 fractionation schemes resulted in an average protein sequence coverage of ∼40%. Likewise, a reduction in HpH fractionation scheme from 46 to 34 and 23 concatenated fractions resulted in nearly identical proteome coverage of 99.4% and 96.7%, respectively (Fig. 5d). This outcome enables the routine acquisition of comprehensive proteomes up to 8 proteomes per day (PPD). The additional proteins identified by multi-shot compared to single-shot analysis are in low abundance range and represent important signaling proteins including transmembrane-receptors and transcription factors (Fig. 5e). Moreover, the fact that our dataset covers ∼80% of core protein complexes in the CORUM database^24,25^, provides evidence of the completeness of our dataset (Fig. 5f). The high peptide coverage also enabled us to search for regulatory post-translational modifications (PTMs), which are often sub-stoichiometric and therefore generally require specific enrichment strategies prior to MS analysis^26^ (Fig. 5g). We identified ∼2700 N-acetylation sites, which as previously shown^2^, mainly target cytoplasmic and nuclear proteins. Additionally, we detected >4000 phosphorylation sites, likely representing the most abundant cellular sites, which in accordance with previous studies^27–29^ are mainly targets of proline-directed kinases such as the cyclin-dependent kinases (CDKs) and the mitogen-activated protein kinases (MAPKs), respectively.

**Figure 5.**
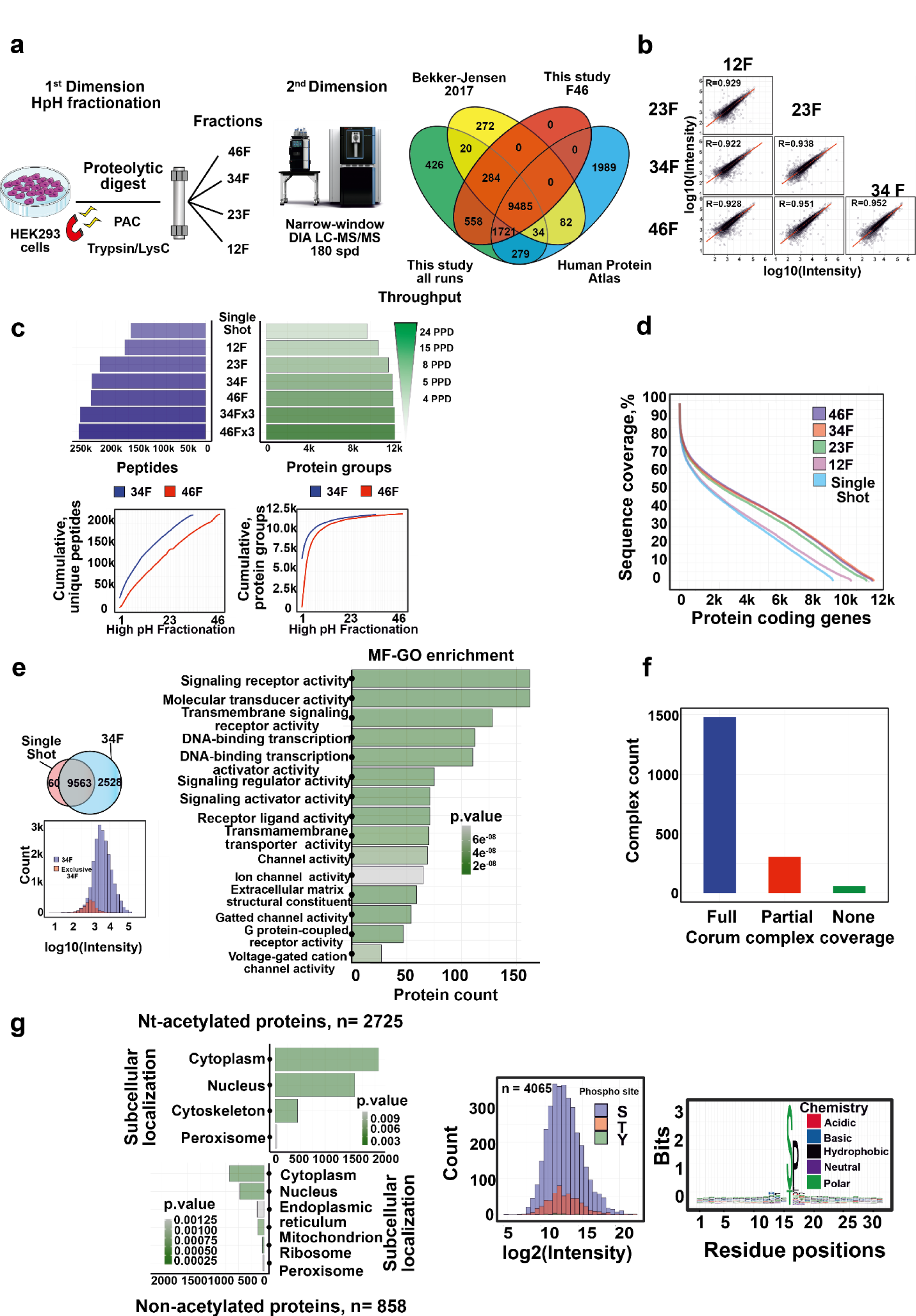
Near-complete human proteomes by multi-shot proteomics. (a) Experimental workflow of all HEK293 experiments and fractionation schemes (left). Overlap of protein-coding genes identified in this HEK293 dataset against the state-of-the-art published dataset and transcriptome (right). (b) Quantitative reproducibility of the method. (c) Protein and peptide identifications for different fractionation schemes and throughput are shown (up). Cumulative peptide identifications by fraction (bottom, left) and cumulative proteins (bottom, right) for both 46 and 34 fractionation schemes are presented. (d) Comparison of sequence coverage achieved by different fractionation schemes and single shot analysis. Overlap of protein groups identified between 34 fractionation scheme (5 PPD) and single shot analysis (2 µg input and 30 min effective chromatographic gradient length).(e) Protein group abundances in HEK293 identified in this dataset compared against single shot analysis (left) is presented. GESEA functional enrichment analysis of the exclusively detected proteins in 34F scheme compared to single shot analysis using gene ontology (GO) molecular function gene set. (f) CORUM protein complex coverage of proteins identified in this dataset. (g) Comparison of the GSEA functional enrichment analysis using gene ontology (GO) cellular component terms gene set of Nt-acetylated and Nt-non acetylated proteome (left). Abundance and sequence logo plot of detected phosphorylation sites without enrichment is shown (right).

### Ultra-fast narrow-window DIA empowers systems biology and clinical proteomics analysis

Functional genomics screens require large-scale experimental approaches to systematically evaluate the functions of genes, often through gene knockout or knockdown^30^. Precise and high-throughput proteomics studies can be used to analyze specific biological processes or diseases caused by a specific genetic perturbation. Historically, the model organism to perform genome-wide screens is *Saccharomyces cerevisiae* and consequently it is a continuous goal of the proteomics community to acquire comprehensive yeast proteomes in the minimum MS analysis time (Fig. 6a). Given the essentially complete yeast proteome profiling provided by the narrow-window DIA strategy with 5-min gradients, we applied this high-throughput strategy to analyze a yeast strain library consisting of 70 gene knockouts targeting cell cycle, proteasome and kinase genes (Fig. 6b). We analyzed the proteome of each knockout strain in biological triplicates and quantified ∼4500 proteins consistently across all conditions with 93.7 % of proteins reproducibly quantified with CV<10% (Fig. 6c). In total, ∼4300 yeast proteins were measured in 80% of the samples (Fig. 6d). Bioinformatics analysis of the knockout-regulated proteins revealed that the observed changes in protein abundances can be attributed to general biological processes such as cellular adaptation of translation rate and metabolic pathways (Fig. 6e). Remarkably, this dataset, collected in a fraction of the time (∼28 h) and at a greater proteome depth for each strain, consistently reproduces the findings of a recent systematic yeast gene knockout screen utilizing proteomics^31^.

**Figure 6.**
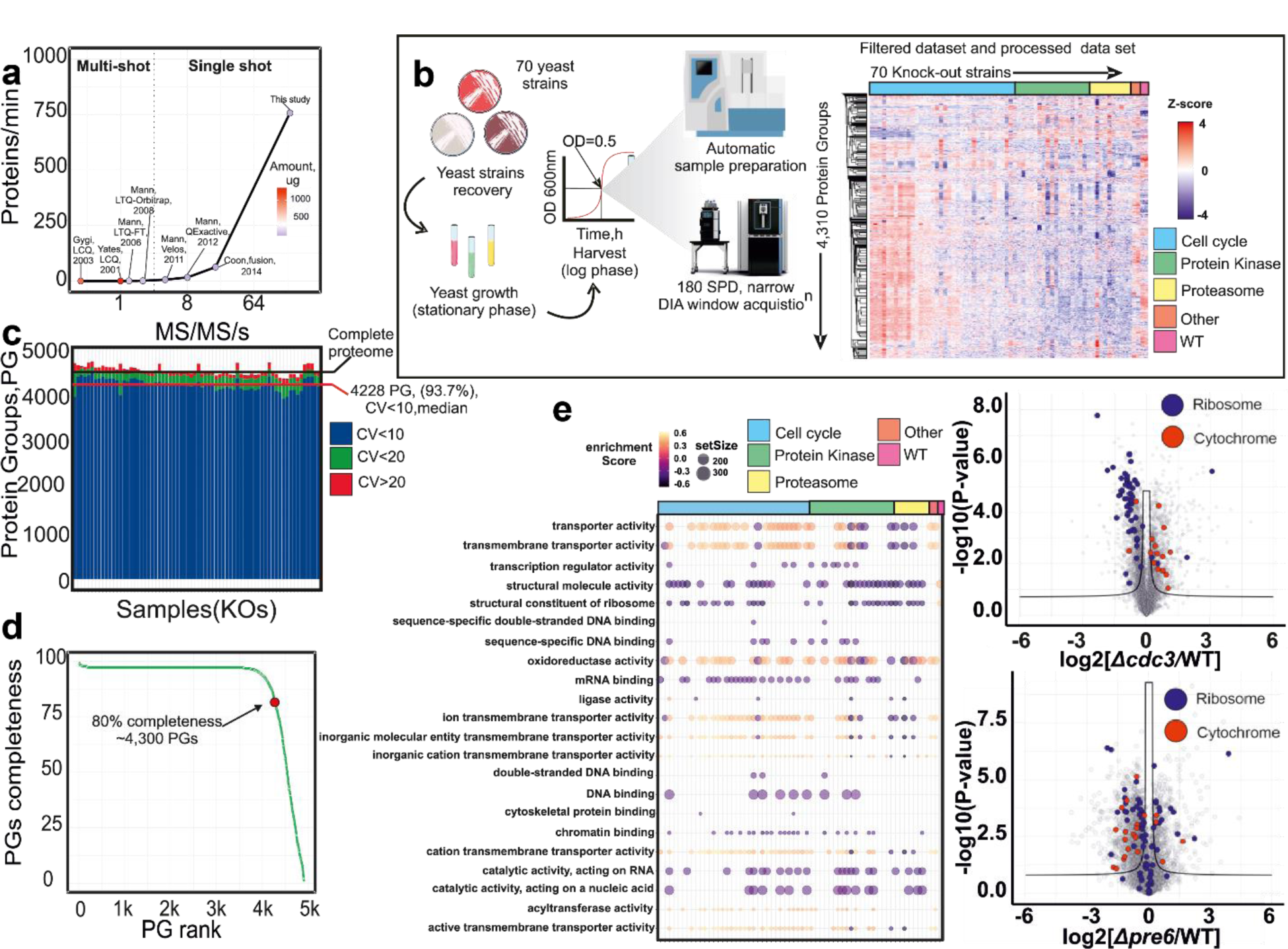
Functional genomics screen by comprehensive yeast proteome profiling. (a) Rate of protein identifications as a function of mass spectrometer MS/MS scan rate for large-scale yeast proteome analysis^15,16,26,32–35^. (b) Experimental setup for high-throughput yeast proteome MS analysis and data set description. (c) Protein identification numbers (mean) and coefficient of variation (CV) per sample (n=3). Data completeness curve showing the number of proteins quantified in a given percentage of samples. GSEA functional enrichment analysis results using gene ontology (GO) molecular function terms gene set. Size of the dot indicates the set size and the color the Normalized Enrichment Score (NES) (left). Volcano plot of differentially expressed proteins between the selected knockouts and a reference strain (right). p-values were obtained by two sided t-test and BH FDR corrected.

Finally, we translated the capability of the narrow-window DIA strategy to a clinical setup using very short 180 SPD gradients to achieve the high-throughput required for large clinical cohorts. As a proof of concept, we performed a fast profiling on the Orbitrap Astral MS of 16 pools of protein extracts from breast capsule biopsies to assess the underlying causes of capsular contracture (n=8 for cases of capsular contracture, and n=8 healthy controls) and the acquired data was processed using Spectronaut v17 directDIA+ method (Fig. 7a). Capsular contracture is one of the most common severe complications to breast implants in patients undergoing breast reconstruction or breast augmentation. The underlying pathophysiology is largely unknown, but it is hypothesized that a low-grade inflammatory response to breast implants leads to a progressively thickening fibrous capsule that contracts around the implant and causes disfigurement and pain. The analysis of all 16 sample pools utilized about two hours of MS time and provided a coverage of 5872 protein-coding genes, with more than 5000 of them quantified in >90% of the samples (Fig. 7b). This fast profiling allowed us to stratify the women diagnosed with capsular contraction by activation of the inflammatory or immune responses (Fig. 7c-e) and highlighted some proteins such as SLC22A4, DEFA1B or MMP7, which could work as biomarkers for capsular contracture (Fig. 7d). Interestingly, MMP7 has been previously identified as a marker for autoimmune inflammation in renal transplant patients^36^.

**Figure. 7.**
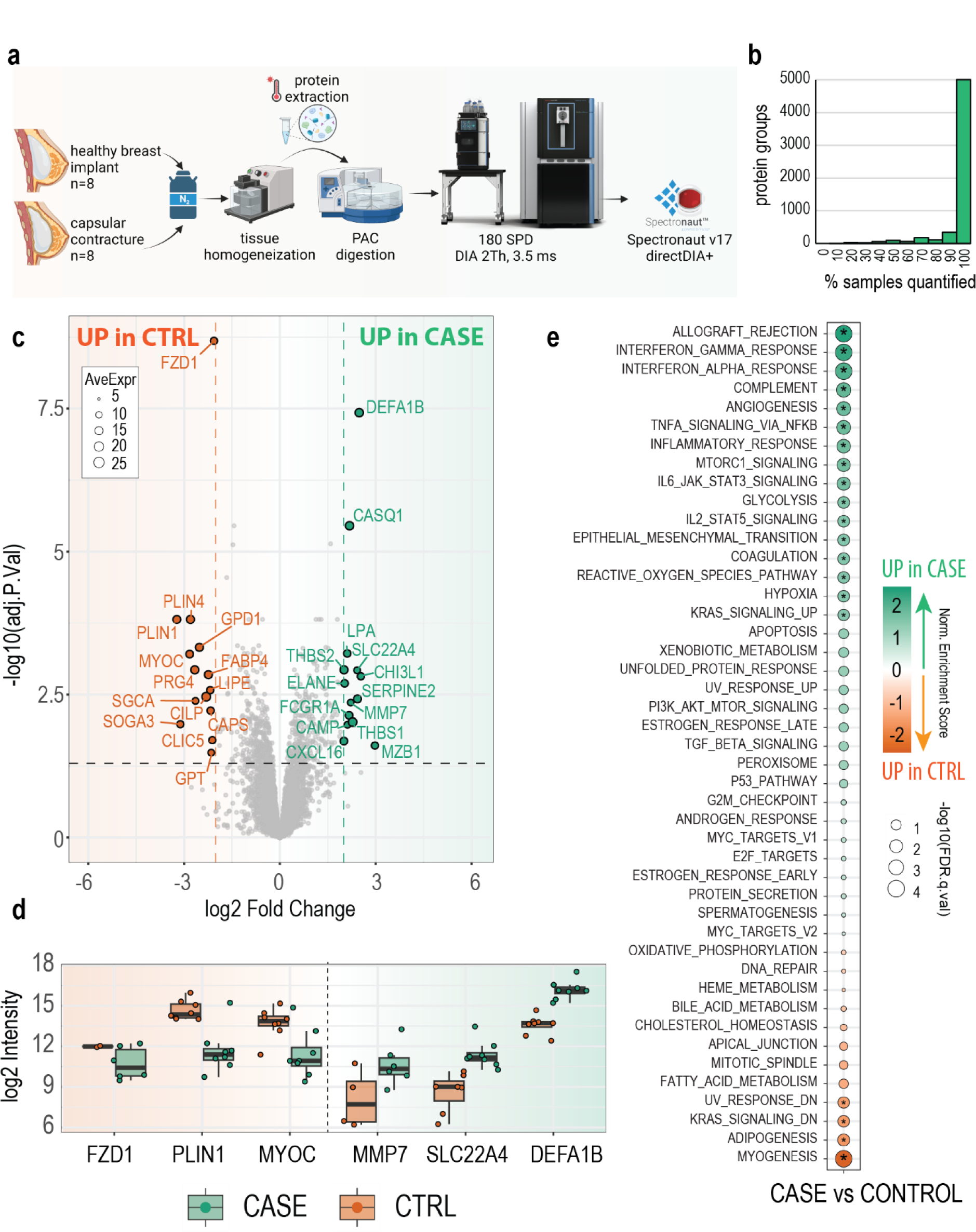
Ultra high-throughput clinical proteomics. (a) Sample preparation workflow of breast implant biopsies for proteome MS analysis. (b) Histogram of data completeness showing the number of proteins quantified in a given percentage of samples. (c) Volcano plot showing differential expression results between control and case samples (limma robust two sample two sided t-test, BH FDR corrected). (d) Boxplot of relevant differential proteins in each patient group. (e) GSEA enrichment analysis results using HALLMARKS gene set. Size of the dot indicates the -log10 of the FDR q-value and the color the Normalized Enrichment Score (NES).

## Discussion

In the last decade, MS-based proteomics has evolved greatly in terms of data acquisition strategies and sample preparation methods, enabling high-throughput analysis of complex proteome samples^7,37–40^. This progress has been primarily driven by technological advances, particularly the introduction of faster scanning mass spectrometers^41,42^.

In this study, we present a different MS acquisition approach based on a novel narrow-window DIA method with an extremely fast scanning Orbitrap Astral mass spectrometer that enables the acquisition of DDA-like DIA data with high proteome coverage and excellent quantitative accuracy and precision with high speed, resolution, and mass accuracy. DIA methods can improve the selectivity by narrowing the isolation window to widths similar to those employed in DDA methods. However, using these settings to monitor the full mass-range was not possible with current state-of-art mass-spectrometers. An example of the utility of narrow-window DIA is high-throughput proteome profiling experiments with short gradients. Here, multiple peptides co-elution increases the prevalence of chimeric spectra, highlighting the importance of using DIA acquisition in this scenario.

Our single-shot data shows that Orbitrap Astral MS provides the sensitivity and acquisition speed needed to routine acquire comprehensive proteomes using narrow-window DIA, maximizing peptide sequencing across the mass range, even with low ion loads. We have optimized the acquisition speed of the Astral analyzer by modulating through the combination of AGC and maxIT, where a 2.5 ms maximum injection time results in >200 Hz MS/MS acquisition rate with 2-Th isolation windows for DIA.

In addition, we showed that narrow-window DIA empowers the routine acquisition of essentially complete human proteomes with a throughput of 5-8 proteomes/day by using very short 5-min LC-MS/MS gradients to analyze the HpH-fractions. This strategy provides ten times higher throughput than previously without any sacrifice in data quality or completeness. The higher peptide coverage resulting from the multi-shot experiment enables direct identification of major PTMs without specific enrichment, providing additional useful data that surpasses the capabilities of next-generation RNA sequencing^2^.

We also demonstrated the capability of the Orbitrap Astral MS narrow-window DIA for comprehensively quantifying multiple proteomes in an unprecedented time scale, which allowed us to map 210 near-complete yeast proteomes in 28 hours.

Altogether, these achievements highlight the potential of MS-based proteomics for quantifying global proteome expression in human samples, such as tumor-derived cell lines and clinical samples.

## Materials and Methods

### Sample preparation

#### Cell lines

Different human cell lines (HeLa, HEK293) were cultured in DMEM (Gibco, Invitrogen), supplemented with 10% fetal bovine serum, 100U/ml penicillin (Invitrogen), 100 μg/ml streptomycin (Invitrogen), at 37 °C, in a humidified incubator with 5% CO2. Cells were harvested at ∼80% confluence by washing twice with PBS (Gibco, Life technologies) and subsequently adding boiling lysis buffer (5% sodium dodecyl sulfate (SDS), 5 mM tris(2-carboxyethyl)phosphine (TCEP), 10 mM chloroacetamide (CAA), 100 mM Tris, pH 8.5) directly to the plate. The cell lysate was collected by scraping the plate and boiled for an additional 10 min followed by micro tip probe sonication (Vibra-Cell VCX130, Sonics, Newtown, CT) for 2 min with pulses of 1s on and 1s off at 80% amplitude. Protein concentration was estimated by BCA.

#### Yeast strains

The *S. cerevisiae* strains used were S288C isogenic yeast strains (MATα) wild-type, BY4743 and BY4743 yeast knockout collection from Horizon Discovery. Briefly, yeast was grown in YPD medium supplemented with G418 at 400 µg/mL at 30 °C. Overnight pre culture was diluted to OD600 = 0.2. Cultured yeast was harvested at OD600 ∼ 0.5 by centrifugation (10,000 x g for 5 min at 4 °C), washed two times with cold PBS and stored at −80 °C until use.

#### Tissue sampling

Samples were obtained from the breast implant capsule of women undergoing exchange or removal of breast implants after breast augmentation. All biopsies were excised from the anterior-inferior part of the fibrous capsule adjacent to the inframammary fold. The biopsies were immediately snap-frozen in a container with dry ice while still in the operating room and stored at −80 °C. Afterwards approximately 100 mg of capsule tissue was transferred frozen to a tissue extraction bag (tissueTUBE TT1, extra thick, Covaris Inc) and placed in liquid nitrogen. The extraction bag was then placed in a Covaris CP02 Automated Pulverizer set to impact level 4 and the instrument was activated to pulverize the frozen sample. This operation was repeated twice. Then the pulverized samples were solubilized with 1000 μL extraction buffer (4% SDS, 100 mM Tris pH 8 supplemented with a cocktail of 1X protease inhibitors (Complete mini, Roche Diagnostics GmbH), 5 mM tris(2-carboxyethyl)phosphine (TCEP), and 10 mM 2-chloroacetamide (CAA), all final concentrations)). Afterwards the samples were sonicated on ice for 1 min (1 sec ON, 1 sec OFF, 50 % amplitude) before heat treatment for 20 min at 80 °C. This was followed by centrifugation at 20,000 xg for 20 min resulting in three layers: A white insoluble pellet, a clear phase in the middle and a milky lipid rich layer in the top. From the clear phase in the middle 3 x 200 μL was carefully transferred to a new tube and this constituted the final protein extract. The remaining lipid rich layer and the pellet were discarded.

#### Preparation of samples for LC-MS/MS analysis

The breast implant capsule tissue samples protein concentrations were determined by the BCA assay. 250 μg of protein were digested by a KingFisher instrument using the protein aggregation capture PAC protocol ^36^. Digestion was performed overnight in 300 μL of 50 mM 4-(2-hydroxyethyl) piperazine-1-ethanesulfonic acid (HEPES) buffer at pH 7.5 supplemented with endoproteinase Lys-C (Wako) and trypsin (Promega) at room temperature. E:P ratio was 1:500 for LysC and 1:250 for trypsin. Sample digests were acidified with tri-fluoro acetic acid to 1% final concentration and stored at −80 °C before pooling and preparation for analysis by mass spectrometry. In total, 48 samples were subject to protein extraction and tryptic digestion as described. In order to keep the samples anonymous, samples representing either cases or controls were pooled three-by-three by adding the same amount of peptide into the pool from each sample contribution to the pool. This resulted in 8 pools with peptides from patients with capsule contracture and 8 pools from controls not seeing contracture.

For global yeast proteome profiling, yeast cells were resuspended 1:2 in lysis buffer composed of 100 mM Triethylammonium bicarbonate (TEAB), pH 8.5, 5 mM, Tris(2-carboxyethyl)phosphine hydrochloride (TCEP), 10 mM chloroacetamide (CAA) and 2% Sodium Dodecyl Sulfate (SDS). Cells were lysed by eight rounds of bead beating (1 min beating, 1 min rest, 66 Hz) in a Precellys 24 homogenizer with 400 µm silica beads (2:1, resuspended cells: silica beads). The extracted protein lysates were heated to 95 °C during 10 min, briefly sonicated and centrifuged at 16,000 g, 4 °C. Afterwards, the protein concentration was approximated using the BCA assay (Pierce^TM^).

The yeast and human cell lines were digested overnight using the PAC protocol^36^. The proteolytic digestion was performed by addition of lysyl endopeptidase (LysC, Wako), and trypsin enzymes 1:500 and 1:250 protein ratio respectively. The samples were incubated at 37 °C overnight. The digestion was quenched by the addition of tri-fluoro acetic acid acid to final concentration of 1%. Peptide mixtures from human cell lines were further concentrated on SepPaks (C18 Vac C18 Cartridge, 1cc/50 mg 55–105 μm Waters, Milford, MA). Final peptide concentration was estimated by measuring absorbance at 280 nm on a NanoDrop 2000C spectrophotometer (Thermo Fisher Scientific). The resulting peptide mixtures from yeast were desalted by Stage-tips, eluted and finally dried down using a speedvac vacuum concentrator. The protein digests from the mixed species for the LFQ analysis were purchased from Pierce™ for HeLa (Cat# 88328), Promega for yeast (V7461), and Waters^TM^ for E.coli (SKU: 186003196). They were mixed manually in six different ratios, E5-H50-Y45, E10-H50-Y40, E20-H50-Y30, E30-H50-Y20, E40-H50-Y10 and E45-H50-Y5, respectively. Samples were kept at −20 °C until further use.

### Offline High pH Reversed-Phase HPLC Fractionation

HEK293 200 µg peptides were separated by HpH reversed-phase chromatography using a reversed-phase Acquity CSH C18 1.7 μm × 1 mm × 150 mm column (Waters) on an UltiMate 3000 high-performance liquid chromatography (HPLC) system (Thermo Fisher Scientific) with the Chromeleon software. The instrument was operated at a flow rate of 30 μl/min with buffer A (5 mM ABC) and buffer B (100% ACN). Peptides were separated by a multi-step gradient as follows: 0–10 min 6.5%B–15%B, 10–59.5 min 15%B–30%B,

59.5–67 min 30%B–65%B, 67–70 min 65%B–80%B, 70–77 min 80%B, 78–87 min 6.5%B. A total of 46 fractions were collected at 60 s intervals. Samples were acidified using 30 µl of 10% formic acid. Samples were dried down using a Speedvac vacuum concentrator. Sample concatenation was performed manually afterwards to the following schemes: 34, 23 and 12 fractions. 200 ng of each sample were injected for LC–MS/MS analysis.

### LC-MS/MS analysis

LC-MSMS analysis was performed on an Orbitrap Astral MS coupled to a Thermo Scientific™ Vanquish™ Neo UHPLC system, and interfaced online using an EASY-Spray™ source. Depending on the gradient used (Table 1) different set-ups were used, either trap-and-elute or direct injection into the column. Column type was chosen also accordingly to the gradient employed (Table 1).

For the DDA experiments, the Orbitrap Astral MS was operated with a fixed cycle time of 0.5 s with a full scan range of 380-980 m/z at a resolution of 180,000. The automatic gain control (AGC) was set to 500%. Precursor ion selection width was kept at 2-Th and peptide fragmentation was achieved by higher-energy collisional dissociation (HCD) (NCE 30%). Fragment ion scans were recorded at a resolution of 80,000, and maximum fill time of 2.5 ms. Dynamic exclusion was enabled and set to 10 s.

For the DIA experiments, the Orbitrap Astral MS was operated at a full MS resolution of 180,000 or 240,000 with a full scan range of 380 − 980 *m/z* when stated. The full MS AGC was set to 500%. Fragment ion scans were recorded at a resolution of 80,000 and maxIT of 2.5 ms. 300 windows of 2-Th scanning from 380-980 m/z were used, unless stated otherwise in Table 2. The isolated ions were fragmented using HCD with 25% NCE.

For the LFQ samples acquired in the Orbitrap Exploris 480 MS, peptides were eluted online from the EvoTip using an EvoSep One system (EvoSep Biosystems) and analyzed in 30SPD (45-min gradient) using a commercial 150 mm analytical column (EV1113 ENDURANCE COLUMN, EvoSep Biosystems). The mass spectrometer was operated in positive mode using DIA mode. Full scan spectra precursor spectra (350–1400 Da) were recorded in profile mode using a resolution of 120,000 at m/z 200, a normalized AGC target of 300%, and a maximum injection time of 45 ms. Fragment spectra were then recorded in profile mode fragmenting 56 consecutive 13 Da windows (1 m/z overlap) covering the mass range 361–1033 Da and using a resolution of 15,000. Isolated precursors were fragmented in the HCD cell using 27% normalized collision energy, a normalized AGC target of 1000%, and a maximum injection time of 22 ms.

Further details are described in Tables 1 and 2.

### Raw MS data analysis

Raw files from DIA and DDA comparison experiments were analyzed in DIA-NN 18.1^43^ using an *in-silico* DIA-NN predicted spectral library (4299848 precursors, allowing for C carbamidomethylation and N-term M excision and 1 missed cleavage). The spectral library was generated from a human reference database (Uniprot 2022 release, 20,588 sequences). The DIA-NN search include the next settings: Protein inference = “Genes”, Neural network classifier = “Single-pass mode”, Quantification strategy = “Robust LC (high precision)”, Cross-run normalization = “RT-dependent”, Library Generation = “IDs, RT and IM Profiling”, and Speed and RAM usage = “Optimal results”. Mass accuracy and MS1 accuracy were set to 0 for automatic. “No share spectra”, “Heuristic protein inference” and “MBR” were unchecked. The MS1 quantification for DDA approach was performed as follows: The output results from DIA-NN were filter for PG.Qvalue < 0.05, Q.Value < 0.01 and Global.PG.Q.Vlaue < 0.01, then DIANN R package was used to calculate the MaxLFQ abundance for protein groups. MaxLFQ abundance was calculated based on “MS1.area” column prior normalization by the ratio Precursor. Normalized/Precursor. Quantity. DIA Approach was assed using MS2-centric methods from DIA-NN output.

Raw files from single-shot dilution series of HEK peptides, window optimization and clinical samples were analyzed in Spectronaut v17 (Biognosys) with a library-free approach (directDIA+) using the human reference database (Uniprot 2022 release, 20,588 sequences) complemented with common contaminants (246 sequences). Cysteine carbamylation was set as a fixed modification, whereas methionine oxidation and protein N-termini acetylation were set as variable modifications. Precursor filtering was set as Q-value, cross run normalization was unchecked. Each experiment was analyzed separately, and those that contained different experimental conditions (different input amounts or acquisition methods) were search enabling method evaluation and indicating the different conditions (each one with n=3 experimental replicates) in the condition setup tab.

Raw files from single-shot analysis of different gradients were analyzed in Spectronaut v18 (Biognosys) using a library-free approach (directDIA+) using the human reference database (Uniprot 2022 release, 20,588 sequences) complemented with common contaminants (246 sequences). Cysteine carbamylation was set as a fixed modification, whereas methionine oxidation and protein N-termini acetylation were set as variable modifications. Precursor filtering was set as Q-value, cross run normalization was unchecked. Each gradient was analyzed separately, and each analysis contained 3 experimental replicates.

Raw files from LFQ analysis of the mixed species samples were analyzed in Spectronaut v17 (Biognosys) using a library-free approach (directDIA+) using a benchmark reference database for the three species (31,657 sequences in total). Cysteine carbamylation was set as a fixed modification, whereas methionine oxidation and protein N-termini acetylation were set as variable modifications. Precursor filtering was set as Q-value, cross run normalization was enabled. Each experiment consisting of samples with same loading amounts was analyzed separately, and each condition contained 3 experimental replicates.

Raw files from different fractionation schemes were analyzed in Spectronaut v17 (Biognosys) with a library-free approach (directDIA+) using the human reference database (Uniprot 2022 release, 20,588 sequences) complemented with common contaminants (246 sequences). Methionine oxidation and Protein N-termini acetylation were set as variable, whereas Cysteine carbamylation was set as fixed modification. Precursor filtering was set as Q-value. Each fractionation scheme was search independently except for searches performed in triplicates. Quantification was performed using MaxLFQ algorithm embedded in iq R package^44^. Briefly, extended Spectronaut output results were filter as follow: PG.Qvalue < 0.01 and EG.Qvalue < 0.01. Then MaxLFQ algorithm was applied using PG. Genes and PG.ProteinNames for protein annotation. Finally, to determine the percent of residues in each identified protein sequence (sequence coverage), the program Protein Coverage Summarizer was used. The human reference database used for quantification and a file containing all detected peptide sequences with a protein name associated (PG.Qvalue < 0.01 and EG.Qvalue < 0.01) were utilized for protein assembling and sequence coverage calculation. For phosphopetides search, 34 fraction scheme was analyzed in triplicates using an empirical library generated *in house* by the HpH fractionation (12 Fractions) of phosphopeptide enrichment (119,793 precursors). Spectronaut output was reformatted using the Perseus plugin peptide collapse^45^ to create a MaxQuant-like site-table.

Yeast Knock-out collection raw files were analyzed using Spectronaut v17 (Biognosys) with a library-free approach (directDIA+) using a database composed of the canonical isoforms of *Saccharomyces cerevisiae* (6059 sequences) complemented with a common contaminant database (246 sequences). Briefly, cysteine carbamylation was set as a fixed modification, whereas methionine oxidation and protein N-termini acetylation were set as variable modifications. Precursor filtering was set as Q-value, cross run normalization was checked.

In Spectronaut (v17 and v18), protein grouping was performed using default protein inference workflow with IDpicker as inference algorithm.

Plots from figures 1b-d were based on MS1 feature detection output retrieved by MaxQuant (v1.6.7.0). Total Ion Current intensity for figure 3c were obtained from MaxQuant (v1.6.14.0). Representative raw files for each method were loaded into MaxQuant and analyzed without the indication of a FASTA file, to only extract the relevant MS features.

## Acknowledgments

We would like to thank Andreas Larsen, Tim Kongsmark Weltz and Mikkel Herly from the Department of Plastic Surgery and Burns Treatment, Copenhagen University Hospital, for access to biopsies from breast implant capsules. Jesper V. Olsen acknowledges funding from the Novo Nordisk Foundation under the Grant number NNF14CC0001. The Novo Nordisk Foundation’s Copenhagen Bioscience PhD Program (grant number NNF16CC0020906) supported U.H.G.

